# Transformer-based modeling of Clonal Selection and Expression Dynamics (TraCSED) reveals resistance signatures in breast cancer

**DOI:** 10.1101/2024.06.15.599136

**Authors:** Nathan Maulding, Jun Zou, Wei Zhou, Ciara Metcalfe, Josh Stuart, Xin Ye, Marc Hafner

## Abstract

Understanding transcriptional heterogeneity in cancer cells and its implication for treatment response is critical to identify how resistance occurs and may be targeted. Such heterogeneity can be captured by *in vitro* studies through clonal barcoding methods. We present TraCSED (Transformer-based modeling of Clonal Selection and Expression Dynamics), a dynamic deep learning approach for modeling clonal selection. Using single-cell gene expression and the fitness of barcoded clones, TraCSED identifies interpretable gene programs and the timepoints at which they are associated with clonal selection. When applied to cells treated with either giredestrant, an estrogen receptor (ER) antagonist and degrader, or palbociclib, a CDK4/6 inhibitor, time-dependent resistance pathways are revealed. For example, ER activity is associated with positive selection around day four under palbociclib treatment and this adaptive response can be suppressed by combining the drugs. Yet, in the combination treatment, one clone still emerged. Clustering based on partial least squares regression found that high baseline expression of both SNHG25 and SNCG genes was the primary marker of positive selection to co-treatment and thus potentially associated with innate resistance – an aspect that traditional differential analysis methods missed. In conclusion, TraCSED enables associating pathways with phenotypes in a time-dependent manner from scRNA-seq data.

## Introduction

While cancer is traditionally considered a disease of genetic alterations, studies have shown adaptive resistance to treatment often involves molecular differences in genetically identical cells (Shaffer, Dunagin, et al.; Rambow et al.; Schuh et al.; Roesch et al.; Sharma et al.; Gupta et al.; Shaffer, Emert, et al.; Emert et al.; Su et al.). The plastic nature of the response to treatments and its heterogeneity may even be a necessity to enable the emergence or selection of resistant clones (Dagogo-Jack and Shaw). Characterizing and understanding tumor heterogeneity and how it affects treatment response is critical to the development of new therapeutic strategies in oncology. Advances in single-cell technologies, such as the tracking of molecular states over time, can be used to address those questions (Emert et al.; Bhang et al.; Biddy et al.; Weinreb et al.; Gutierrez et al.; Oren et al.; Frieda et al.; Umkehrer et al.; Luyi Tian et al.; Leighton et al.; Rodriguez-Fraticelli et al.; Pillai et al.). In particular, transcriptional heterogeneity can be captured through clonal barcoding methods such as TraCe-Seq (Chang et al.). Studies have also shown that “twin” clones (i.e. sister cells with the same barcode) show a higher degree of transcriptional similarity than other clones (Goyal et al.), suggesting that the transcriptional state of barcoded clones persists after multiple divisions. Leveraging this assumption, TraCe-Seq measures the fitness and transcriptional trajectory of clones in parallel pools of cells undergoing different treatments (Figure 1A). To uncover differences associated with the selection process, the fitness of clones at the end of treatment can be mapped back to their transcriptional states pre-treatment or at intermediate time points.

**FIgure 1:**
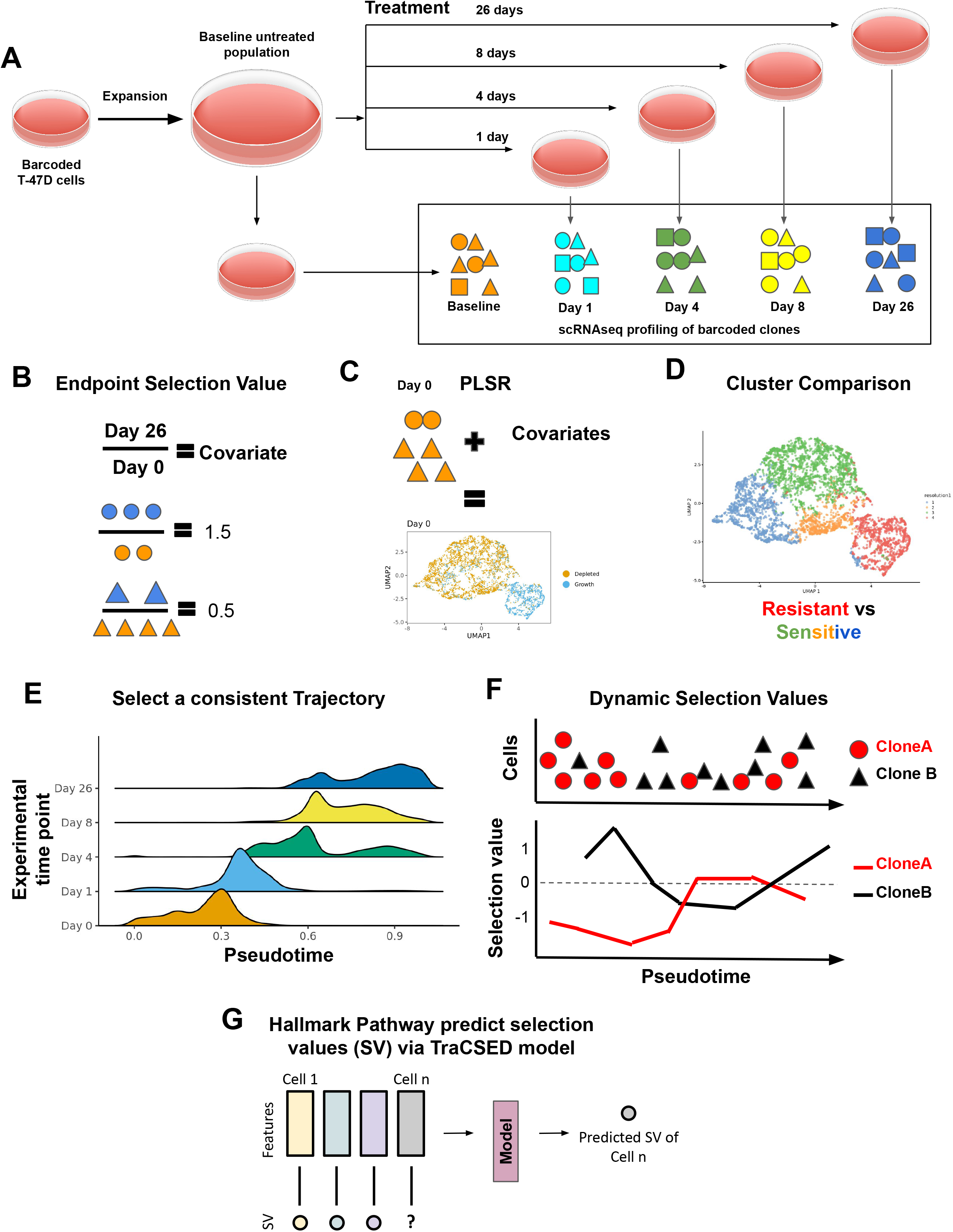
Modeling approaches to study innate and adaptive resistance mechanisms. (A) TraCe-seq experiment includes single-cell RNAseq data from a series of pre-treatment and post-treatment time points that originate from the same pool of barcoded cells. In our case, the treatments include either giredestrant alone (0.1µM), palbociclib alone (0.2µM), or their combination (same concentration). (B-D) Baseline investigations begin by determining the “endpoint selection value” for each clone (circles or triangles) by dividing the fraction of the clone 26 days after treatment by the fraction of clones present pre-treatment relative to the population (B). These covariates are assigned to clones at baseline and combined with the single-cell RNAseq data to create a partial least squares regression (PLSR) dimension reduction (C). A majority of resistant cells are specific to a cluster, which is then compared to clusters with a majority of sensitive cells (D). (E-G) To model adaptive resistance the selection values of clones across the time of treatment must first be determined. This is initiated by selecting a pseudotime cell-by-cell ordering inferred by slingshot which is consistent with the experimental time ordering (E). Clone selection values along the selected pseudotime are determined by dividing the prevalence of the cells ahead of a given cell in the trajectory by the prevalence behind it (F). Finally, clone selection values are paired with gene or pathway features for the single cell to model the prediction of single-cell selection value (G). This prediction considers the temporal context as well as the feature values, such that downstream model interpretability analysis yields dynamic features important in resistance.

While experimental advances have been substantial, methodologies to interpret such multidimensional single-cell transcriptional data sets are still in their infancy. For example, clustering approaches can fail to distinguish resistant from sensitive cell states as recent clonal barcoding systems have shown that a few genes, or even a single gene, can determine a cell’s fate (Richman et al.). However, incorporating clonal information through a tunable parameter into the dimensionality reduction has been shown to benefit the identification of features associated with cell outcomes (Richman et al.). Innate resistance may play a major role as intrinsic cell states conferring resistance could be encoded prior to external stimuli (Goyal et al.). Additional forms of non-genetic response may also contribute to resistance. For example, altering epigenetics or rewiring signaling networks could underlie adaptive resistance that emerges after treatment has begun (Cerezo et al.). Because traditional methods rarely integrate longitudinal data as a continuum and associate them with an outcome, they yield limited understanding of adaptive resistance, its underlying gene programs, and the time at which such changes occur.

To address those shortcomings, we first developed a semi-supervised approach with partial least squares regression (PLSR) to identify pre-treatment markers of innate resistance in TraCe-seq datasets (Figure 1A-D). This approach incorporates the transcriptional similarities of cells together with a phenotypic response to reveal signatures that might otherwise be hidden from traditional methods. PLSR uncovers single gene markers of innate resistance in breast cancer, such as SNHG25 and CLDN1, which were missed by PCA-based clustering and differential expression methods.

Next, to identify gene programs associated with adaptive resistance, we introduce TraCSED (Transformer-based modeling of Clonal Selection and Expression Dynamics). TraCSED learns a dynamical process of clonal selection using a clone trajectory’s inferred pseudotime from TraCe-Seq single-cell RNA-seq data as the time series input for model fitting (Figure 1E-G). It uncovers interpretable gene programs that are directly related to the selection process and estimates time periods at which these may be critical for resistance.

We applied PLSR and TraCSED to infer resistance mechanisms for two treatments used in the clinic to treat breast cancer, palbociclib and giredestrant, which target CDK4/6 and, respectively, estrogen receptor (ER). We compared the results of single-agent treatments to combination therapy in which we observed different factors associated with resistance. In particular, all clones that resisted the single-agent treatments became sensitive to the combination except one that was still resistant to the combination. This demonstrates the value in TraCSED in finding important pathways associated with a phenotypic variable, in this case clone fitness, and how this information can guide our biological understanding of drug response.

## Results

### Treatments induce different clonal selection and transcriptional responses

We performed a TraCe-seq experiment with T-47D cells treated with giredestrant, a selective estrogen receptor (ER) antagonist and degrader, or palbociclib, a CDK4/6 inhibitor, for 1, 4, 8, or 26 days (Figure 1A). The acute response from T-47D cells to either drugs is cytostatic as shown by the normalized growth rate (GR) values close to zero (Supplementary Figure 1A) (Hafner et al.). Both treatments showed a decrease of clone diversity, quantified by a reduction in the number of unique clonal barcodes in the cell population (Figure 2A). We classified clones as either negatively selected (NS) or positively selected (PS) based on the ratio of clonal frequency in the overall population at day 26 versus day 0, which we termed the “endpoint selection value”, a surrogate for clone fitness (Figure 1B). Clones with a greater frequency after 26 days of treatment would have an endpoint selection value greater than 1 and would be considered PS, whereas clones with less frequency would be less than 1 and NS. As an illustration, when we selected the 11 clones with at least 200 cells across all time points, we observed differences as to which clones were PS for giredestrant compared to palbociclib treatment (Figure 2B).

**Figure 2:**
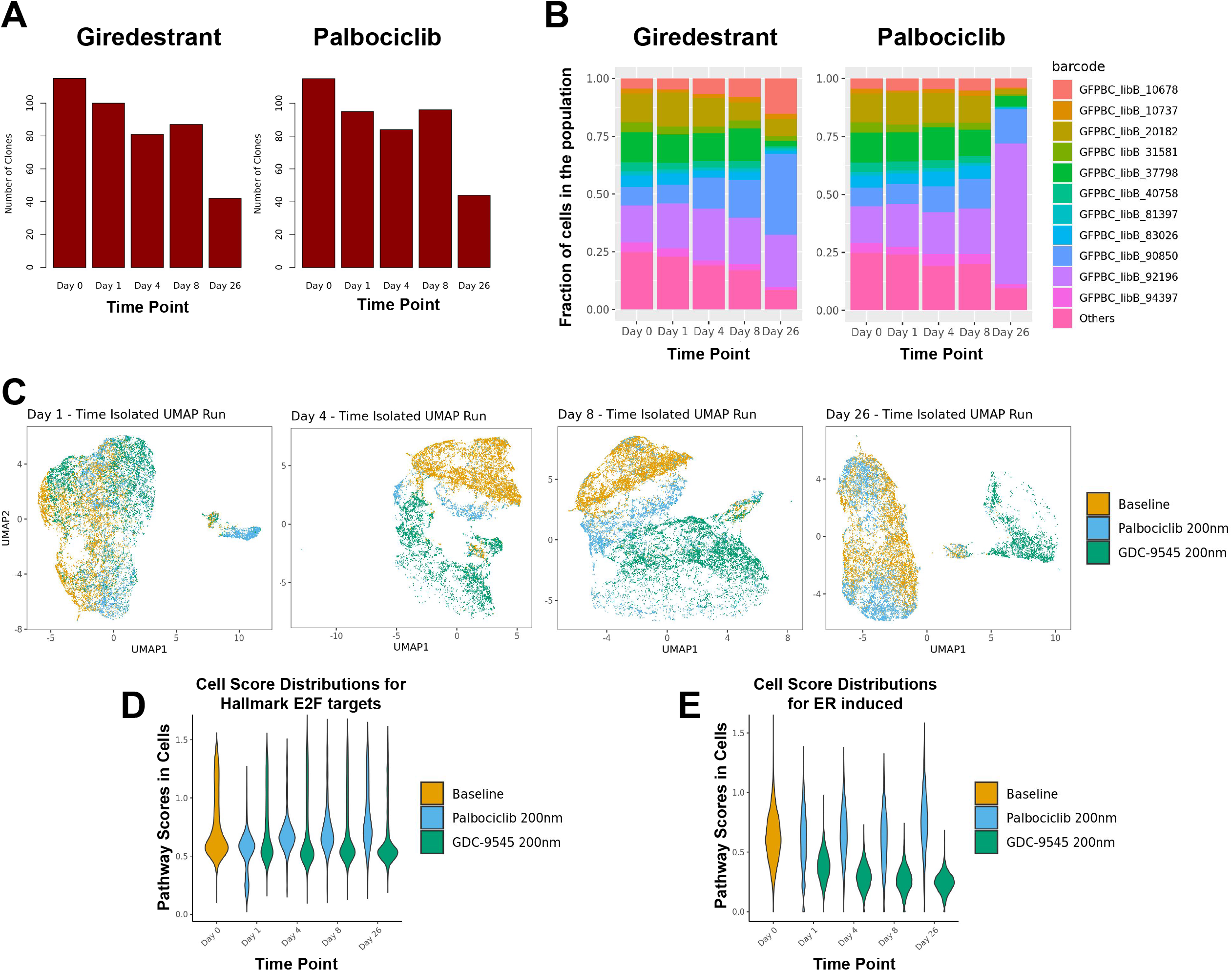
Giredestrant and palbociclib induce unique clonal selection and transcriptional responses over time. (A) Overall number of clones detected in samples of T-47D cells treated with either giredestrant (left) or palbociclib (right) at different time points. (B) Fraction of cells for clones with a minimum of 200 cells for giredestrant (left) or palbociclib (right) treatments at different time points. (C) UMAP of cells from the baseline sample and each time point individually for both giredestrant or palbociclib treatments. (D-E) Pathway scores from scuttle for the E2F targets pathway (D) and ER activity (E) for baseline cells and cells treated with either palbociclib or giredestrant at different time points.

UMAP based on PCA showed that both treatments induced a strong transcriptional response with clear separation of the cells in each treatment group, suggesting the dynamics of response are characteristic of each treatment (Figure 2C, S1B). By day 26, the separation in transcriptomes between treatments was clear with cells from the palbociclib-treated sample overlapping in UMAP space with baseline cells. In contrast, the giredestrant-treated cells formed a separate UMAP cluster, reflecting a more pronounced shift in the transcriptional state. In terms of pathway scores, reduction of ER activity is sustained throughout the time points of the giredestrant-treated samples (Figure 2E), but the fraction of cells with E2F positive score returns to baseline levels in both giredestrant- and palbociclib-treated samples (Figure 2D). Giredestrant’s inhibition of ER activity was maximal around 4-8 days aligned with its effect on cell cycle (Figure 2E). Consistent with this result, we observed by Western Blot a downregulation of ER and its target, the progesterone receptor (PR) in the giredestrant-treated sample (Supplementary Figure 1C) and a regain of pRb, a marker of proliferation, in both PS populations. The PS populations for each treatment were, as expected, more resistant than the parental population to the treatment used for selection as shown by the positive GR values reflecting partial growth inhibition (Supplementary Figure 1A, C). PS cells can proliferate, albeit slower when rechallenged with palbociclib treatment, whereas the PS cells in the giredestrant treatment became completely insensitive to giredestrant. We also observed a partial cross-resistance between the two drugs (Supplementary Figure 1A): the palbociclib-PS population had much GR values for the response to giredestrant than the parental population and the reverse was also true.

### PLSR identifies baseline signatures associated with selection of clones

The observed “rebound” in the E2F proliferation signature score could be due either to an innate fitness advantage of some clones prior to treatment, which allow those clones to outcompete other ones, or it could be due to an adaptive response induced by the treatment itself in the PS clones. In either case, we hypothesized that the initial transcriptional state influences the likelihood of a clone being positively selected under treatment. However, initial attempts using pseudobulk samples in which cells from clones of similar endpoint selection values were grouped revealed few differentially expressed genes for either treatment condition (Supplementary Figure 2A). Alternatively, unsupervised clustering via PCA did not separate PS and NS clones for either treatment due to high gene expression variability across cells of one clone compared to the separation achieved through PCA clustering (Supplementary Figure 2B). Thus, both pseudobulk differential expression (DE) analysis and PCA-based clusters failed to identify informative resistance pathways.

We therefore undertook a more sensitive approach to detect resistance pathways using semi-supervised partial least-squares regression (PLSR) dimensionality reduction (see Methods; Figure 1C). PLSR incorporates the transcriptional state as independent variables and the endpoint selection value as a dependent variable to search for a data projection in which PS and NS clones are well separated in the lower dimensional space. Using the PLSR-based UMAP, cells from PS clones were better separated from cells from NS clones (Figure 3A,B). In this projection, we were easily able to define clusters that were strongly enriched clones that were positively selected across time (Supplementary Figure 2C, 2D) revealing baseline markers of selection (Supplementary Figure 2E).

**Figure 3:**
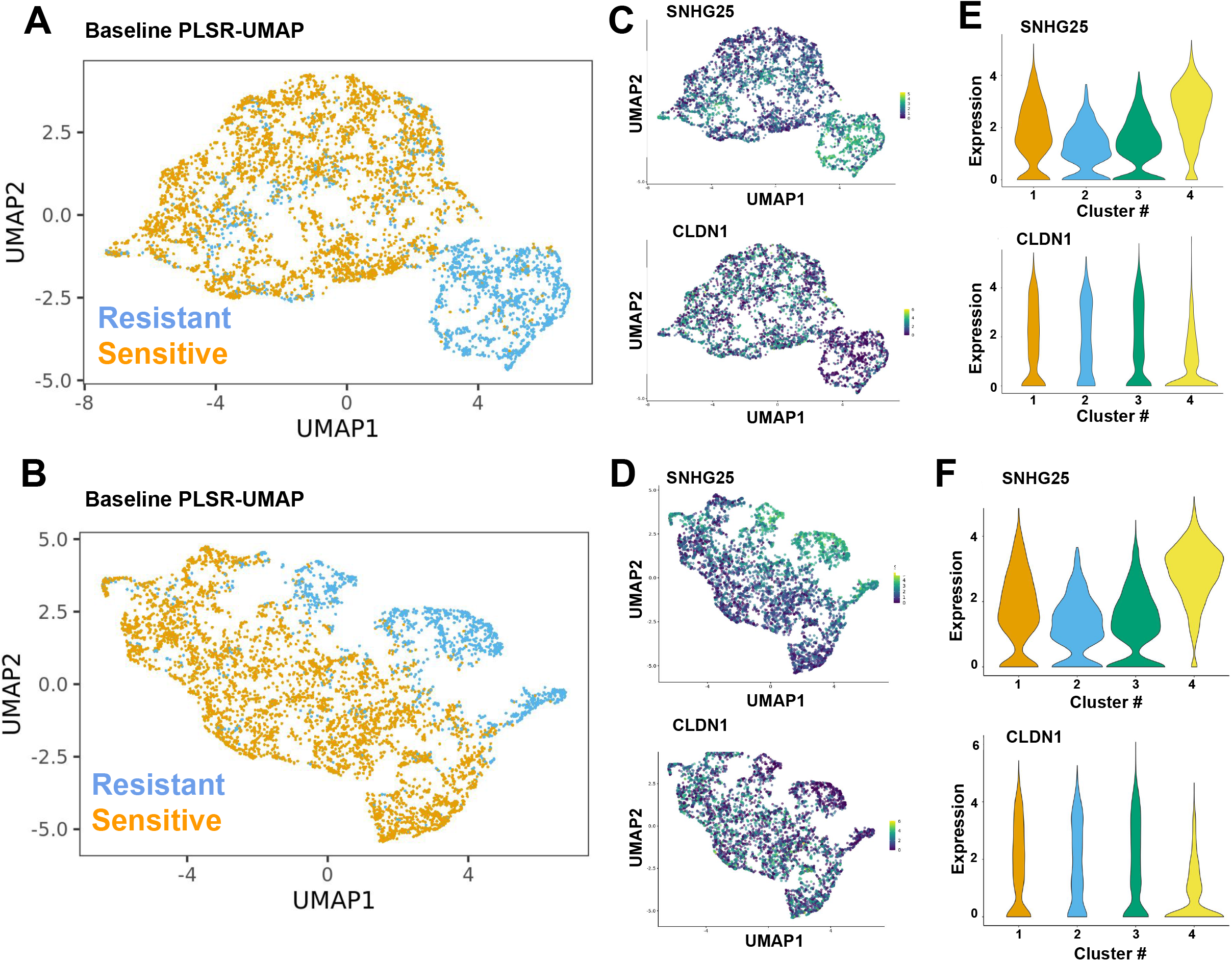
Partial least squares regression (PLSR) reveals baseline signatures hidden by traditional approaches like differential expression. (A-B) PLSR-based UMAP for baseline cells associated with outcome of either giredestrant (A) or palbociclib (B) treatment. Positively selected (PS) cells are in blue and negatively selected (NS) in orange. (C-F) SNHG25 and CLDN1 expression levels are displayed in the UMAP (C-D) and violin plots (E-F) for giredestrant (C,E) or palbociclib (D,F). Cluster 4 contains most PS cells for both giredestrant and palbociclib (see Supplementary Figure 1D).

PS cells were aggregated into a single PLSR cluster for giredestrant treatment (Figure 3A), which enabled a pseudobulk differential gene expression analysis comparing expression observed in cells of the PS cluster to cells outside the cluster. Among the differentially expressed genes (DEGs), high SNHG25 and low CLDN1 expression were identified (Figure 3C, 3E). Similarly, a PS cluster was also identified for the palbociclib-treated cells (Figure 3B), with SNHG25 and CLDN1 identified as DEGs for the palbociclib PS cluster (Figure 3D, 3F). CLDN1 is a known marker used for classifying subtypes of breast cancer, and low CLDN1 is a marker for aggressive triple-negative BRCA and predictive of recurrence (Zhou et al.; Morohashi et al.; Lu et al.; Fougner et al.). SNHG25 has been shown to be associated with tumor activity, but is poorly understood (Liu et al.; Zhiyu et al.; He et al.; Zeng et al.; Biagioni et al.).

### TraCSED fits the adaptive resistance using a generative model

As both positively and negatively selected clones had a strong transcriptional response as illustrated in the UMAP (Fig S3C-E), we extended our modeling framework to leverage the longitudinal transcriptional data and explore potential acquired resistance mechanisms. We therefore took a dynamical modeling approach to identify the transcriptional programs associated with the clonal selection process revealed by the TraCE-seq data. We assumed that a population of cells ordered along a continuous cell trajectory captured the progression from sensitivity to resistance of individual clones. We used the Slingshot method to estimate the cell trajectory and project the cells along a response curve, establishing a pseudotime ordering of the cells. We calculated selection values across the Slingshot pseudotime for each clone (Figure 4A-B, see Methods). To associate transcriptional changes with treatment response, we developed an interpretable generative model that predicts selection values based on a set of pathway features, which we called TraCSED for Transformer-based modeling of Clonal Selection and Expression Dynamics. After the Slingshot pseudotime is established, we calculated the selection value along the trajectory for every cell belonging to a clone (Figure 1E-F, Figure 4A-B). The selection value for a cell belonging to clone *c* at time *t* was calculated as the fraction of clone *c*’s cells ahead of time *t* in the pseudotime ordering divided by the fraction of clone *c*’s cells behind time *t*. The transformer was then trained to predict the selection value of the cell at time *t* using the transformer’s features as well as the features and selection values of clone *c* cells in the context of the cell of interest (Figure 1G). Testing of model accuracy occurred over four pseudotime intervals (Figure 4C). In this way, the model learned a conditional relationship between the features and selection values based on where the cell resided in pseudotime.

**Figure 4:**
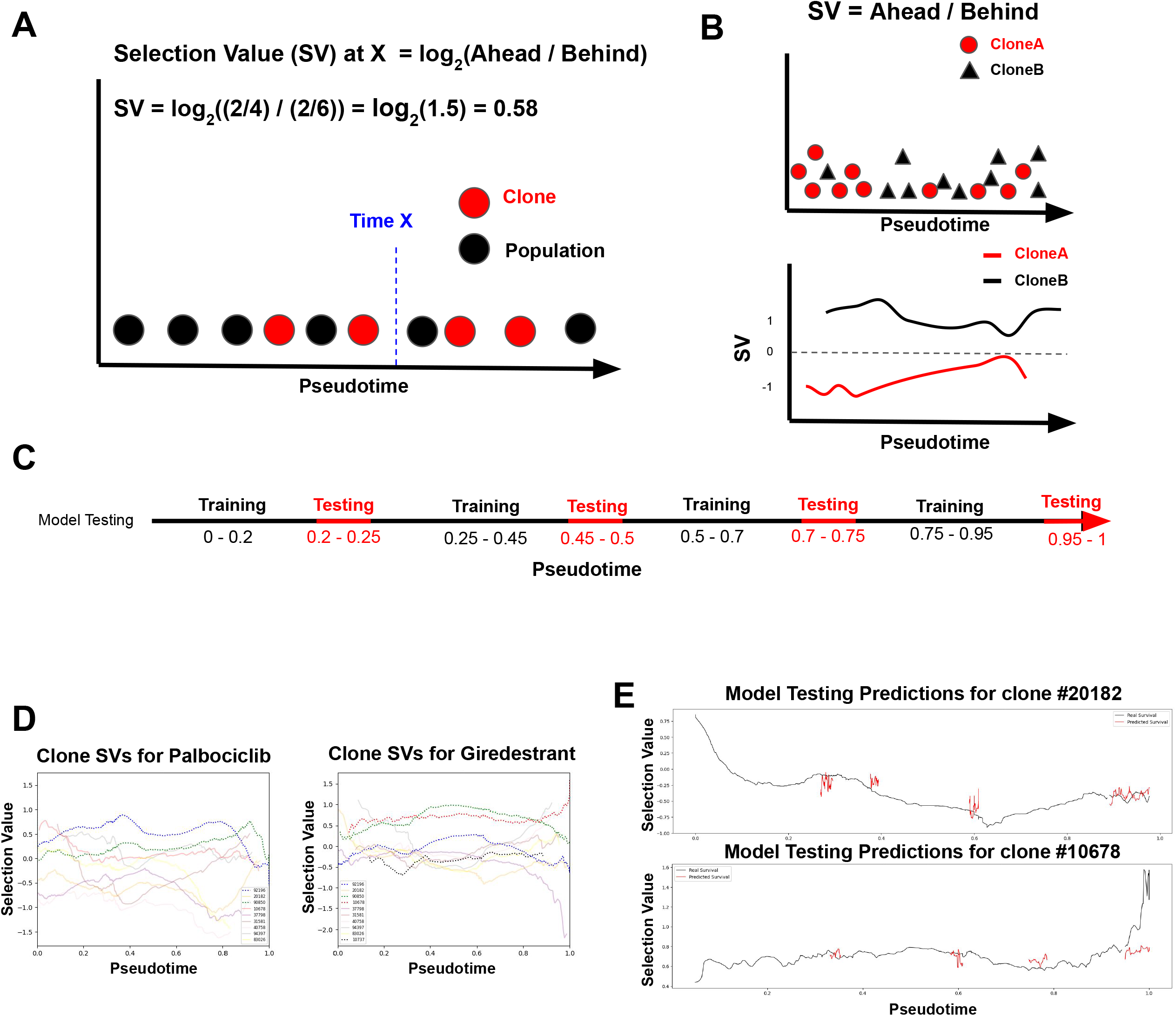
Transformer-based modeling of Clonal Selection and Expression Dynamics (TraCSED) predicts selection values across pseudotime. (A-C) An example clone (red) exists as part of a total population of cells (black) across a pseudotime progression. To determine the selection value at any given time (blue) for the clone in question the fraction of cells ahead and behind a given pseudotime is used (A, see Methods). This results in a continuous selection value curve for each clone across pseudotime (B). This model of dynamic selection is then tested in four different increments (red) along pseudotime to ensure a quality model fit at all time points (C). (D) The resulting transformation from raw data along pseudotime to selection value curves for each clone is shown for each clone in palbociclib (left) and giredestrant (right) treatments. (E) Example of the model testing (red) for two clones under giredestrant treatment is displayed.

We required a minimum number of cells (set to n=200) to be present across the time course to model the selection process of a specific clone as selection values were too noisy otherwise (see Methods). 11 clones in total fit this criteria for at least one treatment (Palbociclib - 9, Giredestrant - 10, Combination treatment - 11). In this case, we used the individual cell Hallmark pathways scores (Liberzon et al.) as features of the model. Across all testing periods (see Methods), the model has lower error than a multivariate regression (Supplementary Figure 4), but the MSE values varied between testing periods. In general, the prediction was worse for the last testing interval as the selection value was noisier due to its dependence on the number of cells, which decreased at the end of the trajectory (Figure 4E). When assessing the contribution of the attention and convolutional layers, we found that the combination of both sets of layers yielded the best compromise of fit quality and prediction accuracy (Supplementary Figure 4). Thus, TraCSED can fit interpretable features associated with the positive or negative selection process of specific clones.

### Model pathway features reveal adaptive resistance mechanisms

Permutation of TraCSED’s input layer enables the identification of features that induce changes to the predicted selection value, a procedure called permutation importance (Altmann et al.). In addition, by altering the inputs as a function of pseudotime, a dynamic pathway importance score can be calculated, enabling the interpretability across time for individual clones. Using this procedure, we found that permuting scores from “Epithelial Mesenchymal Transition” (EMT) resulted in a deviation from the true selection value (upper panel) and a spike in feature importance (middle panel) within a pseudotime range that aligns with the 4 to 8 day time points of giredestrant treatment (lower panel) (Figure 5A). This exemplifies how the model is made interpretable for a single clone. In order to understand whether these selection mechanisms observed were more universal, we plotted pathway scores and feature importance for all modeled clones together. Scores for the EMT pathway in PS clones were decreased compared to NS clones between 0.5-0.9 pseudotime for giredestrant treatment (Figure 5B).

**Figure 5:**
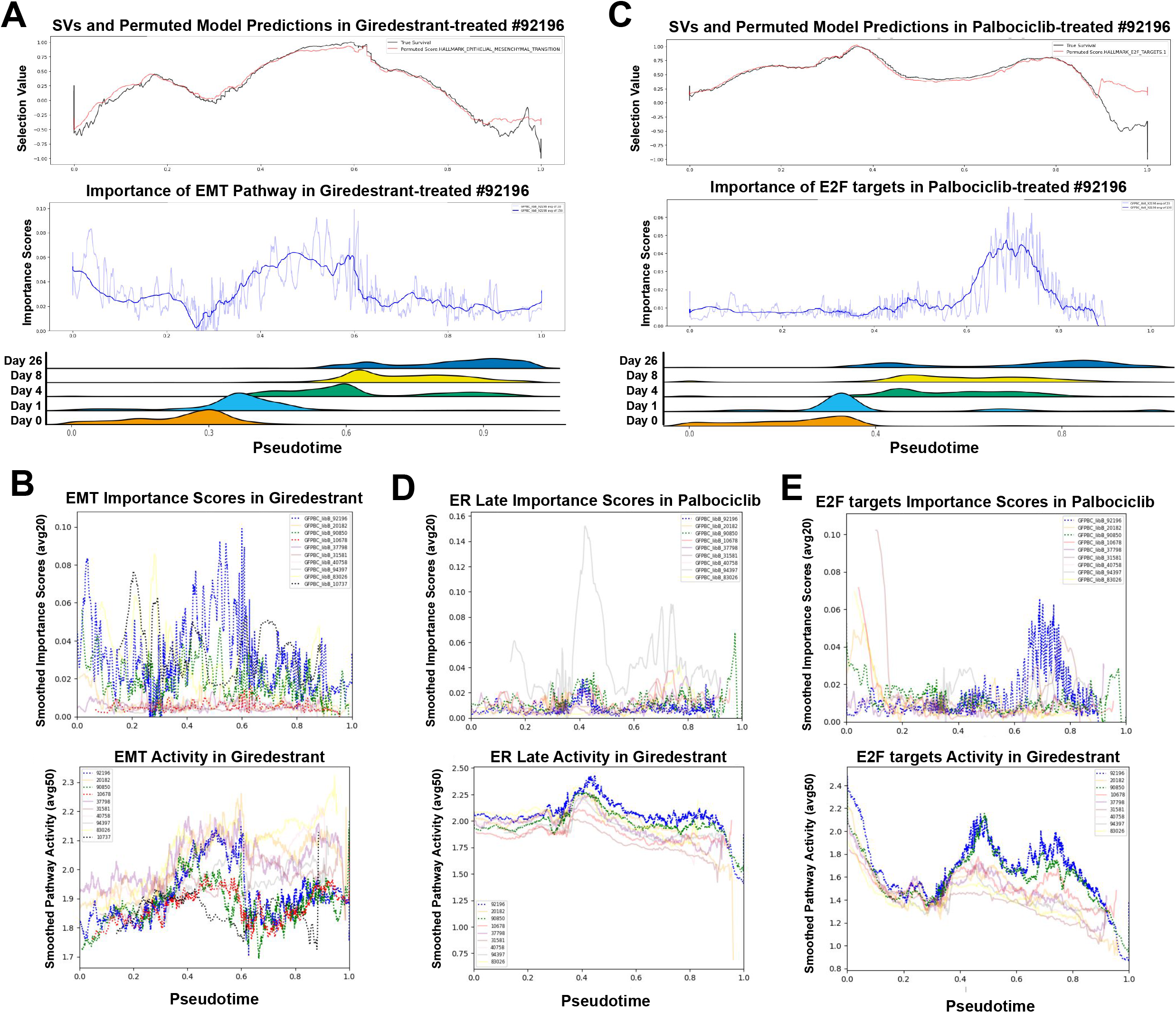
TraCSED identifies pathways associated with PS clones. (A) Model predictions with permuted EMT activity (red) were plotted against the true selection values (black) under giredestrant treatment (A, top) for a specific clone. (Middle) Feature importance for the EMT pathway with averaging smoothing of 20 (light blue) or 150 (dark blue) adjacent cells along pseudotime. (Bottom) Ridge plot of all cell distributions for each experimental time point along pseudotime. (B) Smoothed model importance (averaging 20 adjacent cells) was then plotted for PS clones (bold, dashed lines) and NS clones (faint, solid lines) in the giredestant treatment for the EMT pathway (top). Smoothed (averaging 50 adjacent cells) EMT pathway activity (bottom). (C-E) Same a A-B for palbociclib treatment. True selection values (black) and model with permuted E2F targets (red) (C, top); feature importance scores for E2F pathway (C, middle); and ridge plots of cell distributions for palbociclib treatment (C, bottom). Smoothed model importance (top) and smoothed pathway activity (bottom) plotted for Estrogen Response Late (D) and E2F targets (E) in palbociclib treatment.

Similarly, we examined the feature importance for clones undergoing palbociclib treatment. For example, the E2F target score was important between 8 to 26 days (Figure 5C). When looking at all clones together, the “Estrogen Response Late” scores for PS clones were important and increased above NS clones in pseudotime ranges of 0.35-0.5 corresponding to the early time points (Figure 5D). This Estrogen Response signature in PS clones preceded another signature where E2F target scores increased within pseudotime ranges of 0.6-0.8 (Figure 5E). Other pathways were similarly noted as important in selection including “G2M checkpoint” in palbociclib and the AKT pathway in giredestrant (Supplementary Figure 5). Taken together, these findings suggested that single-agent treatment was not sufficient to overcome adaptive resistance but the combination of palbociclib and giredestrant might be effective because the giredestrant may prevent the positive selection of clones in the palbociclib treatment that had higher ER activity.

### Adaptive resistance is reduced with combination treatment

Next, we treated cells with palbociclib and giredestrant as a combination to investigate our earlier finding. Combination therapy proved to be effective in suppressing proliferation and ER activity across the whole population, which was not previously achieved by palbociclib or giredestrant alone (Supplementary Figure 6A-B). While most clones that were PS in either single agent treatments became NS in the presence of combination, one clone was PS in both single-agent treatments and remained PS in the combination treatment after six months (Figure 6A). The selected population is indeed much more resistant to treatment than individual resistant populations (Supplementary Figure 1B). Similar to the giredestrant-PS population, the surviving population lost ER and PR expression compared to the parental population suggesting independence on ER activity (Supplementary Figure 6C). In addition, surviving cells have lower cyclin D expression, while retaining high pRB, showing less dependency on CDK4/6-CyclinD activity to proliferate (Supplementary Figure 6C). Our PLSR method identified two different clusters associated with PS (Figure 6B and S6D). Similarly to individual treatments, high SNHG25 was also associated with resistance in combination, but in contrast, CLDN1 no longer showed enrichment in the PS clusters. Additionally, SNCG was specifically associated with resistance in combination therapy (Figure 6C-E). SNGC is a synuclein protein known as a marker for late-stage breast cancer (Wu et al.; Lu Tian et al.; Zhuang et al.), implying that cells resistant to combination treatment could be reflecting a cellular state found in tumors from late-stage patients.

**Figure 6:**
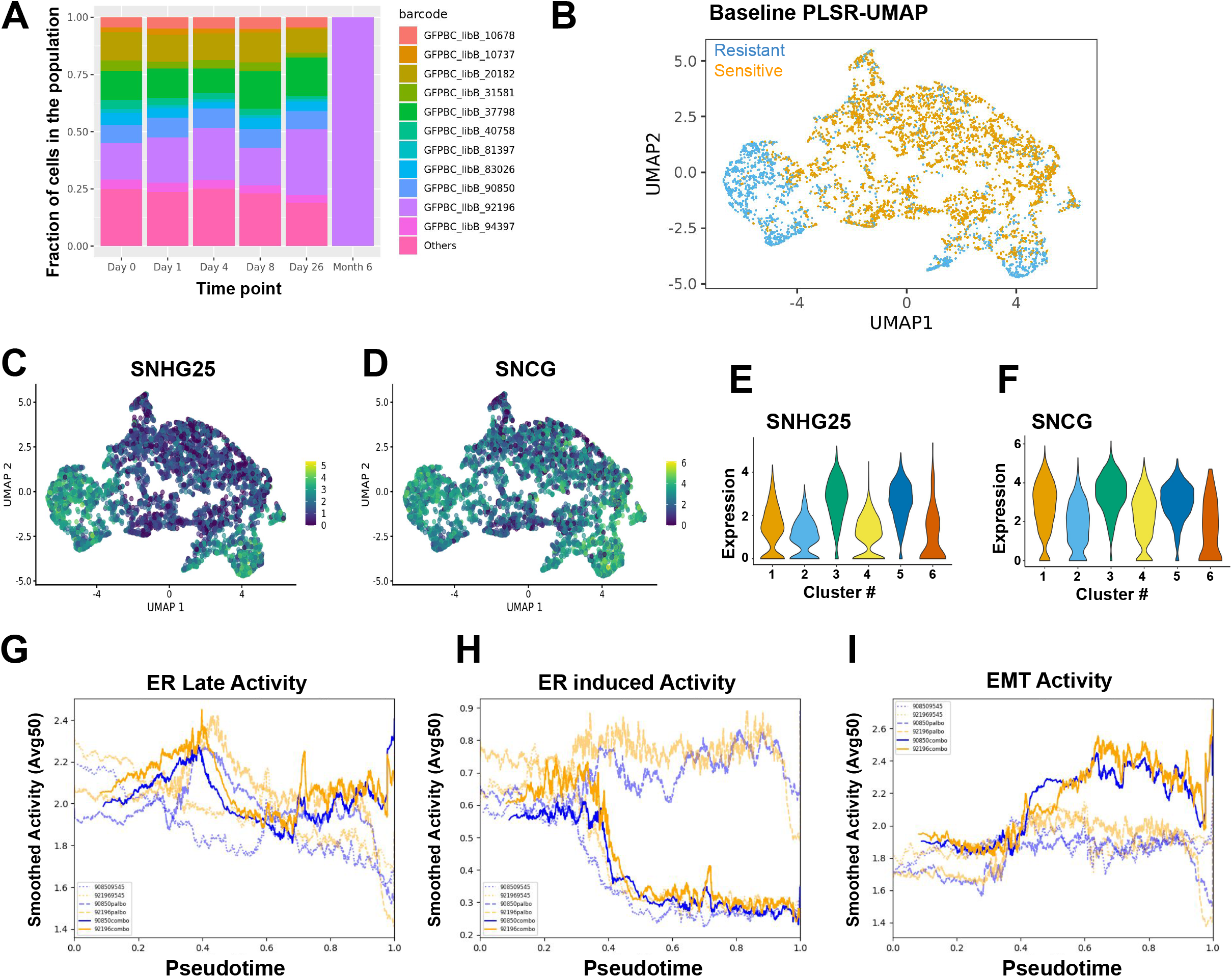
Combination of palbociclib and giredestrant reduces the importance of adaptive resistance mechanisms. (A) Fraction of cells for clones with a minimum of 200 cells in the combination of giredestrant and palbociclib at different time points. (B) PLSR-based UMAP for baseline cells associated with outcome combination treatment. Positively selected (PS) cells are in blue and negatively selected (NS) in orange. (C-D) Gene expression in baseline cells was shown via UMAP from (B) for SNHG25 (C) and SNCG (D). (E-F) Violin plot of SNHG25 (E) and SNCG expression (F) in baseline cells. PS cells are in clusters 3 and 5 (see Supplementary Figure 6C). (G-I) Pathway activity for a PS clone (orange) and a stable clone (blue) plotted for palbociclib (faint, dashed lines), giredestrant (faint, dotted lines), and their combination (bold, solid lines) using gene sets for Estrogen Response Late (G), ER induced (H), and Epithelial Mesenchymal Transition activity (I).

We then used TraCSED to characterize the dynamic selection process occuring in the combination treatment. Because the Estrogen Response signature was important in the selection process for palbociclib alone and preceding higher E2F targets signature scores, we were curious to see the effect of suppressing ER signaling with giredestrant. We observed that both ER and proliferative signatures had lower activity and showed reduced importance in predicting the selection values of PS clones (Supplementary Figure 6E-F) compared to what was observed for the palbociclib treatment. Similarly, the EMT pathway had low activity in PS clones under giredestrant treatment, but in the combination treatment all clones had similar levels of EMT pathway activity (Supplementary Figure 6G).

While the combination treatment is overall more potent, one PS clone (#92196) still emerged after 6 months. Scores for key gene signatures such as “Hallmark Estrogen Response Late” (Figure 6F), “ER induced” (Figure 6G), or “EMT” (Figure 6H) pathways were not substantially different than a clone (#90850) that is outcompeted at 6 months. The lack of differentiation is consistent with our model. It indeed identified that the features associated with positive selection in either single-agent treatment are substantially less important in the combination treatment compared to the single-agent ones. Therefore, high baseline expression of the SNHG25 and SNCG genes found through PLSR are the primary markers of positive selection for combination treatment and differentiate clone #92196 from the other ones (Supplementary Figure 7). In conclusion, TraCSED is complementary to PLSR for identifying the different factors associated with adaptive and innate resistance. PLSR highlighted markers of innate resistance that are consistent with late-stage cancer phenotypes, whereas TraCSED allowed us to identify vulnerabilities in adaptive resistance to single agent treatments that were addressed through combination treatment.

## Discussion

Giredestrant and palbociclib were effective in blocking proliferation of T-47D cells and resulted in a selective bottleneck as reflected by the reduction of the number of clones over time. This process was associated with changes in the transcriptomes of cells, in particular down-regulation of ER and, respectively, proliferative signaling. However, some clones were positively selected and differences were observed in the selection process between the two single-agent treatments (Figure 2A-E). This positive selection can occur due to an innate resistance detectable in pre-treatment conditions, or it could be a result of an emerging adaptive resistance. To investigate these mechanisms, we developed a PLSR approach for finding markers of innate resistance (Figure 1B-D), and a transformer-based model, called TraCSED, for finding markers of dynamic selection in adaptive resistance (Figure 1E-G).

We found that both unsupervised clustering and differential expression analysis comparing PS and NS clones of pseudobulked samples to study innate resistance was not effective in this study (Fig S2A-B). On the other hand, supervised clustering based on PLSR balanced intrinsic variability of the clone with phenotypic/ outcome and enabled us to isolate PS clones into a single cluster for giredestrant (Figure 3A) and palbociclib (Figure 3B), which revealed markers unique to the PS cluster (Supplementary Figure 2E). Two examples of these PS markers were SNHG25 and CLDN1. In giredestrant and palbociclib treatments, SNHG25 is increased in the PS cluster and CLDN1 is decreased (Figure 3C-F). SNHG25 is associated with a variety of cancers (Liu et al.; Zhiyu et al.; He et al.; Zeng et al.), and is part of a family of long-noncoding RNAs (SNHGs) with multi-functional roles in cancer progression, particularly in the EMT pathway (Biagioni et al.). In addition, CLDN1 levels are well-documented for their role in EMT and breast cancer, have been used previously for subtyping different types of cancer such as the claudin-low basal subtype in breast cancer (Herschkowitz et al.), and correlate with breast cancer recurrence (Zhou et al.; Morohashi et al.; Lu et al.; Fougner et al.). PCA-based clustering was not able to identify those genes (Supplementary Figure 2F), and neither of these PS markers were found with pseudobulk differential expression (Supplementary Figure 2F), demonstrating the value of PLSR in finding mechanisms of innate resistance.

In addition to innate mechanisms of resistance, we observed transcriptional changes in both PS and NS clones after treatment (Supplementary Figure 3C-E). Because existing methods do not associate outcomes like selection with temporal transitions, we developed TraCSED, a dynamic, context-aware and interpretable model composed of attention and convolution layers, to investigate mechanisms of adaptive resistance. The dynamic selection for each clone is modeled across pseudotime (Figure 1E-G; Figure 4A-B) and then tested at four pseudotime intervals (Figure 4C, E). The TraCSED model facilitates interpretability by highlighting features important for the selection process of individual clones (see Methods). The EMT pathway had high model importance and low activity for PS clones in giredestrant treatment (Figure 5B), suggesting this pathway may be associated with resistance. In palbociclib treatment, there was high model importance and increased activity in the Estrogen Response Late pathway (Figure 5D) which preceded a subsequent increase in the E2F targets pathway (Figure 5E). Thus in both cases, TraCSED captured an adaptive resistance mechanism to single-agent treatments.

When combining giredestrant and palbociclib, only one PS clone survived 6 months of combination treatment (Figure 6A). Consistent with our hypothesis, the adaptive resistance pathways we identified with TraCSED in the single-agent treatments were no longer associated with dynamic selection in the combination (Supplementary Figure 6C-E). Additionally, the single surviving PS clone did not demonstrate activity in the ER and EMT pathways differing from NS clones (Figure 6F-H). Thus, only the high SNHG25 and high SNCG levels (Figure 6C-E and S7) identified by the PLSR approach in the two PS clusters differentiated the PS clone from the NS ones suggesting that innate resistance mechanisms were driving resistance to giredestrant and palbociclib combination treatment. Increased SNCG is a known marker of resistance and late-stage Breast cancer (Wu et al.; Lu Tian et al.; Zhuang et al.), suggesting that the innate cellular state of these PS clusters may resemble a state found in late-stage tumors. While we elected to use pathway features for our modeling for compute efficiency, increased interpretability, and reduced noise, our modeling approach can also be applied at the gene level. In cases where data quality allows for gene-level analyses, TraCSED may be able to identify individual genes associated with resistance that may not be captured in pathway-level analyses.

Determining how a cell will respond to treatment and how it might be resistant is an ongoing field of study. Here we developed a PLSR approach and the TraCSED model for determining markers of innate and, respectively, adaptive resistance. With PLSR, we were able to identify SNHG25 and CLDN1 markers of positive selection in single-agent treatment, which were not found by traditional methods of pseudobulk and unsupervised clustering. Through TraCSED dynamic modeling we found pathways associated with adaptive resistance, including increased ER activity in palbociclib treatment. We were then able to show that the adaptive resistance is reduced when palbociclib is combined with the ER suppressing activity of giredestrant. Consistently with our hypothesis, our model identified that no feature was associated with an adaptive resistance in the combination treatment, leaving the innate resistance marker, SNHG25 and SNCG, found with the PLSR model, as the main features associated with positively selected cells. Altogether, our two complementary modeling approaches contribute to study and characterize non-genetic sources of therapeutic resistance.

## Methods

### Datasets and Dependencies

T-47D breast cancer cells were labeled with TraCe-seq library as previously described (Chang et al.). A subculture was established from 300 cells to establish the input cell population. These 400 cells were expanded to 2 million cells total, and randomly distributed to the following treatment groups in replicates: 200 nM giredestrant treatment, 200 nM palbociclib treatment, and 200 nM giredestrant + 200 nM palbociclib combination treatment. Cells were trypsinized after 1, 4, 8, or 26 days of treatment and profiled by single cell RNA-sequencing using Chromium Single Cell 3’ Reagent Kits (10x Genomics). Baseline scRNA-seq profiling was obtained after 1 day of treatment in DMSO. In addition, for the combination treatment, cells were harvested and profiled after 6 months following exposure to both compounds.

### Drug response

T-47D (female, adenocarcinoma from pleura) ER+ breast cancer cell lines were cultured using standard aseptic tissue culture techniques at 37°C in RPMI medium supplemented with 10% FBS, 2mM L-Glutamine (Catalog 10440, Sigma), 1X Minimum Essential Media, Non-Essential Amino Acids (MEM NEAA, Catalog No. 11140-0500, Thermo Fisher) and 1X Antibiotic-Antimycotic (Catalog No.15240-112, Thermo Fisher). Cell line ancestry was determined using Short Tandem Repeat (STR) profiling using the Promega PowerPlex 16 System. Cells were treated for 7 days with either 0.2µM of giredestrant, 0.2µM of palbociclib, or their combination and the relative number of viable cells was determined by CyQUANT (ThermoFisher, C7026). Growth rate normalization was performed (Hafner et al.).

### Single-cell analysis

For pathway features, we used Hallmark pathways (Liberzon et al.) and a select set of curated pathways. Single-cells were scored on their pathway activity using the scuttle package in R (McCarthy et al.). Activity across time and treatments is shown via violin plot. Additionally, PCA dimensionality reduction to 5 principal components, followed by UMAP plotting was used to view how cells transcriptomic features change throughout time in the different treatments.

### Quality control

The number of clones present across time and treatment was plotted. Clones with 200 cells or more were displayed with a stacked barplot showing the various clones’ relative proportion across time and treatment (Figure 2B). Clones were considered positively selected if the proportion of the cells increased relative to the population. All clones that did not have a total of 200 cells across time were combined and displayed as “other”. No bias in read counts between PS and NS clones or across treatments and time points was observed.

### Partial least squares regression (PLSR)

Phenotypic comparison based on pseudobulk was performed with DESeq2 differential expression (DE) (Love et al.) on clonal response to treatment in which clones that increased over their baseline percentage by 26 days of treatment (resistant) were compared to those that were absent after 26 days (sensitive). Additionally, expression based PCA clustering was performed to group cells by similar transcriptomes.

For the partial least squares regression (PLSR) method (Abdi), the endpoint selection value is included as a covariate for the dimensionality reduction of expression. The endpoint selection value is a ratio of the cells present after 26 days of treatment over the cells present at baseline. The first 5 components are then used to plot a UMAP representation of the data. Leiden clustering was then done at various resolutions. Pairwise distance of PS and NS cells was performed for both the PLSR and PCA methods to compare which is better at distinguishing the two states. To understand how the clones are distributed across the PLSR clusters, a heatmap displaying the fraction of the clone population in each cluster was plotted. Alongside the heatmap is included the increase (red) or decrease (blue) for each clone compared to pretreatment conditions for Days 1, 4, 8, and 26 after treatment (Supplementary Figure 2D). Resistant clones were concentrated in a single cluster which was then compared to sensitive clusters via the findMarkers package by scran (Lun et al.). PS (blue) and NS cells (orange) were plotted as a UMAP. PS markers found through findMarkers were plotted via UMAP and violin plots.

### Overview of the dynamic generative model

The fundamental steps of the generative modeling approach are 1) selecting an appropriate pseudotime trajectory representation of the data, 2) determining the selection values for each clone across pseudotime, 3) modeling the selection values of each clone, 4) evaluating the model for overfitting, and 5) assessing feature importance across time through permutations of the model. This method allows for the prediction of selection values, but its primary focus is to avoid overfitting and to create feature interpretability across time.

### Selecting an appropriate trajectory representation

In order to model the single-cell selection value of a particular clone, cells must be ordered along a continuous trajectory with start and end points consistent with the experimental design. In this case the trajectory continuum is made using Slingshot pseudotime inference (Street et al.). Selecting a representation that closely resembles the experiment is done through taking the TRACE-Seq data and 1) performing dimensionality reduction through PCA or PLSR, 2) producing Leiden clustered representations at various resolutions, 3) performing Slingshot inference with assigned start and end clusters and normalizing pseudotime, 4) selecting an appropriate pseudotime representation through a summation of KS statistics between TRACE-Seq time points. This results in a pseudotime representation that can be used for modeling clonal selection values.

The TraCe-Seq data is represented in both an unsupervised (PCA) and semi-supervised (PLSR) fashion for comparison. Leiden clustering is then performed on a series of resolutions between 0.2 and 1. Start and end clusters are then selected (based on consistency with the experiment and maximizing distance between start and end points) as input for each and used as input for Slingshot. The pseudotime inference produced by slingshot is then normalized on a 0-1 scale, where the first percentile of pseudotime is set as the minimum value, 0, and the ninety-ninth percentile of pseudotime is set as the maximum value, 1. Each of these normalized pseudotime series is then assessed via a KS statistic summation (Massey), where the pseudotime distribution of Day 0 is compared to Day 1, Day 1 to Day 4, etc. The trajectory representation with the largest KS statistic summation is then selected and used for modeling clonal selection values.

### Determining clonal selection values for modeling

Modeling resistance in an interpretable way begins with clonal selection values. With four different testing periods and a quality model fit being the utmost priority for interpretability, clones must have a minimum of 200 cells across time to be modeled. This left 11 clones across treatments which have sufficient cells. To make the selection value a dynamic metric, we map it across the pseudotime continuum. This is done by considering the number of cells of the clone of interest compared to the total population. At each pseudotime *t* that a cell exists, a calculation is made where the cell population “ahead” of the cell of interest in pseudotime is divided by the cell population “behind” it (Figure 4A) using the following equation:

> log_2_((# cloneA after t / # population after t) / (# cloneA before t / # population before t))

To avoid the strong fluctuations occurring at the start and end points of the selection value curves due to a smaller number of cells present at the extremes (Figure 4B), we fit a smoothing process through iteratively averaging 5 adjacent cells (Figure 4D). Then, a boundary threshold is placed to preserve the regions of pseudotime with less than 0.15 max-normalized absolute change in selection value.

### Framework and modeling design choices

Our primary objective is an interpretable model across pseudotime that avoids overfitting, whereas extrapolation beyond the experimental time is not of interest. Since the modeling is sequential and we want to evaluate the fit across the whole pseudotime, we decided to test our model on multiple intervals instead of only testing the end of the pseudotime. We withheld four intervals that represented 20% of the pseudotime for testing: [0.2-0.25], [0.45-0.5], [0.7-0.75], and [0.95-1] (Figure 4C). Additionally, clones are trained and tested separate from each other to create individual models. This facilitates interpretability and keeps resistance mechanisms distinctive between clones.

### Evaluation of the model

In order to evaluate the contributions of different aspects of the model, we employed three “control” approaches: regression, a transformer without convolutional layers, and a transformer without attention layers. Regression provides a simple baseline by which to assess whether the temporal context adds value to the model fit. Subtracting the convolution or attention aspects of the model, reveals the influence of these aspects’ importance to the architecture for generalizing an accurate learning rule from the data. These models are then assessed via mean squared error (MSE) on the testing data in the four time periods described above (Figure 4E and S4).

### Temporal importance of features through permutation

To assess the influence of features on the prediction of the selection value we used permutation feature importance (Altmann et al.). In this approach we randomized a feature of interest 30 times and used the model with the altered feature to make a prediction. Changes in an important feature result in increased prediction error from baseline. For the 30 permutations of a feature, we compute the error at each pseudotime position to assess which features are important in predicting the selection value at specific pseudotimes. The mean error at each position is used in downstream analysis.

### Interpretability of the model

As an example of how feature permutation provides interpretability to the model of dynamic selection in a particular clone, the true selection value curve was plotted (black) with the permuted model’s selection value prediction (Figures 5A and 5C). Below this was the model importance scores for the permuted pathway with various smoothing factors (averaging the adjacent 20 or 150 scores). Ridge plots of the pseudotime distributions of real time were plotted to understand how the pathways found to be important can be attributed to periods within the experiment.

Mean error scores from each of the clone models were smoothed by averaging the adjacent 20 scores and plotted together. The activity for the feature is also plotted along pseudotime with a similar smoothing process of averaging the adjacent 50 scores. In each of these plots, PS clones are shown in bold, dashed lines and NS clones in faint, solid lines (Figures 5B and 5D-E). This was done for the Epithelial Mesenchymal Transition, Estrogen Response Late, and E2F targets pathways. To compare palbociclib, giredestrant, and combination treatment the activity scores for a PS clone and a stable clone were plotted for EMT and ER pathways. In these plots the pathway activity of the PS clone (orange) is plotted for each of the three treatments across time with the activity of the stable clone (blue) so that the two clones and treatments can be assessed (Figure 6G-I).

## Software Availability

The code used for the TraCSED model and PLSR investigation of innate resistance in this paper can be found at https://github.com/Genentech/TraCSED.

## Data Availability

Single-cell data can be accessed in GEO under the series GSE260703 (reviewer access token:).

## Acknowledgements

We would like to thank the Roche Advanced Analytics Network for funding which supported this work. Also, the TraCe-Seq working group and developers from Genentech for the building of the technology and for providing helpful feedback.

## Author Information

NM, JS, XY, and MH designed the study; JZ, WZ, and XY performed the experiments; NM and MH performed the analyses; CM, JS, XY, and MH supervised the study; NM, CM, JS, XY, and MH wrote the manuscript.

---

**Figure S1.**
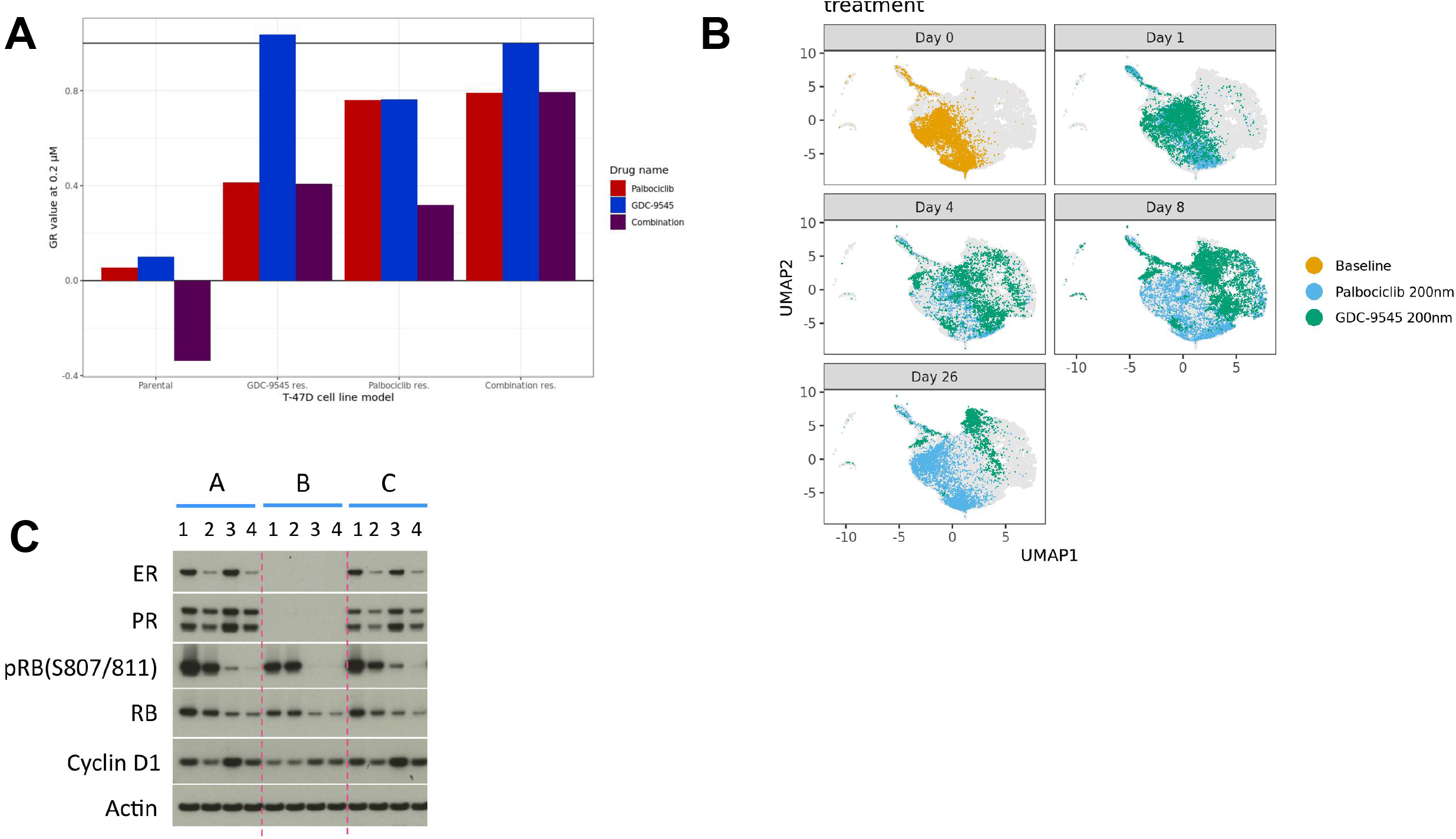
(A) GR value for parental T-47D, giredestrant-treated PS population, or palbociclib-treated PS population, or combination-treated PS population treated for 7 days with either 0.2µM of giredestrant, 0.2µM of palbociclib, or their combination. (B) Transcriptomic changes are observed in palbociclib treated cells (blue), but return to a baseline (orange) state, while giredestrant-treated cells (green) have remained changed. (C) Western Blot of parental T-47D [A], giredestrant-treated PS population [B], or palbociclib-treated PS population [C], treated for 24 hours with either DMSO [1], giredestrant [2], palbociclib [3], or their combination [4].

**Figure S2.**
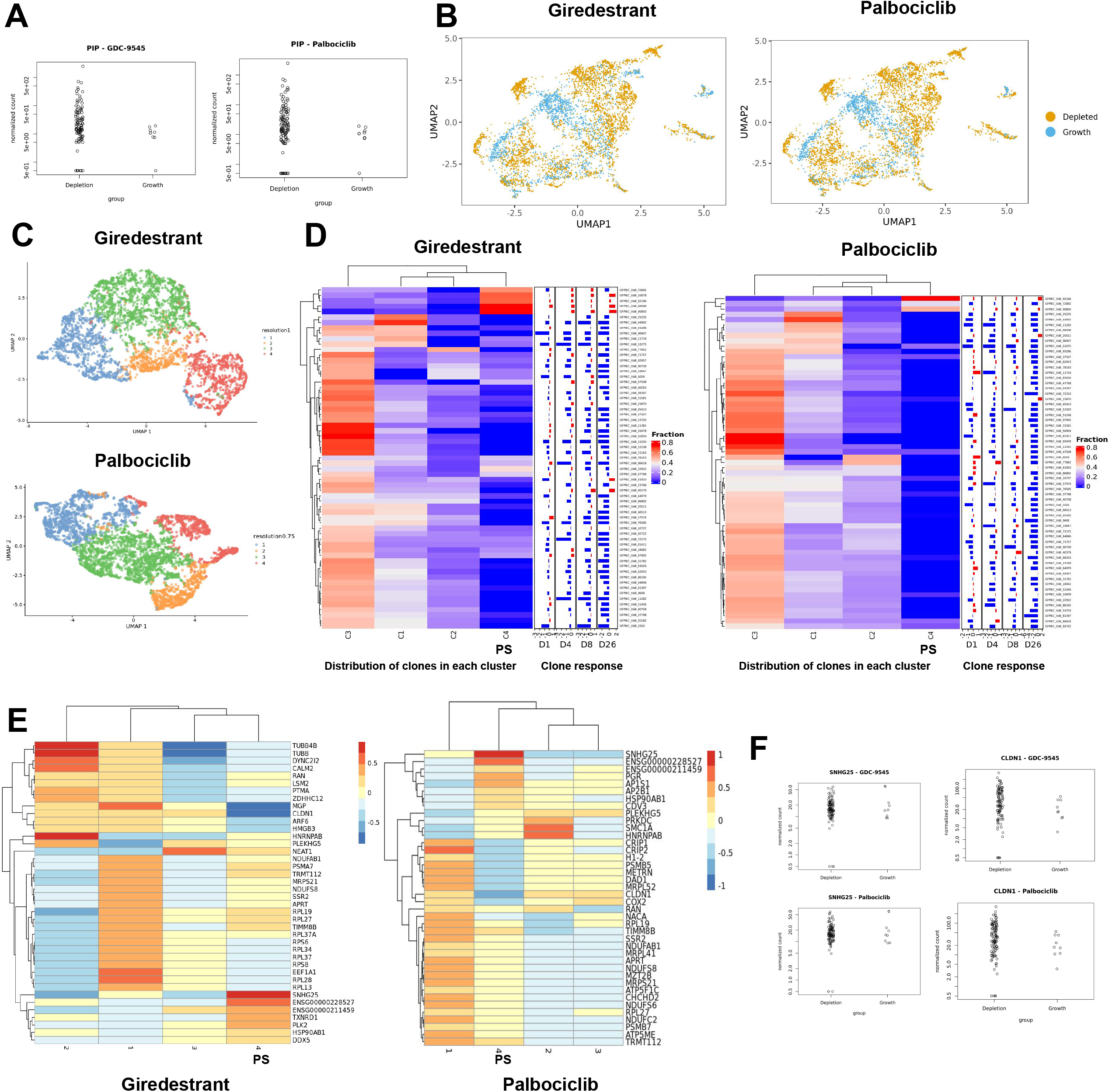
The PIP gene is differentially expressed in PS clones compared to NS clones in giredestrant and palbociclib (A). PS clones (blue) are not separated from NS clones (orange) when PCA dimension reduction is used to create a UMAP projection (B). Clustering results produced from the PLSR approach are plotted for giredestrant and palbociclib-treated cells (C). Heatmaps depict the fraction of clones (y-axis) within a given PLSR cluster (x-axis), where cluster 4 is the PS cluster in both giredestrant and palbociclib treatment. The barplots on the right side show the increase (in red) or decrease (in blue) in the clonal population at days 1, 4, 8, and 26 after treatment (D). Genes found to be differentially expressed via findMarkers are plotted relative to the expression in the PS cluster 4 (E). Genes found through the PLSR approach, SNHG25 and CLDN1, were not shown to be differentially expressed when comparing PS clones to NS clones (F).

**Figure S3.**
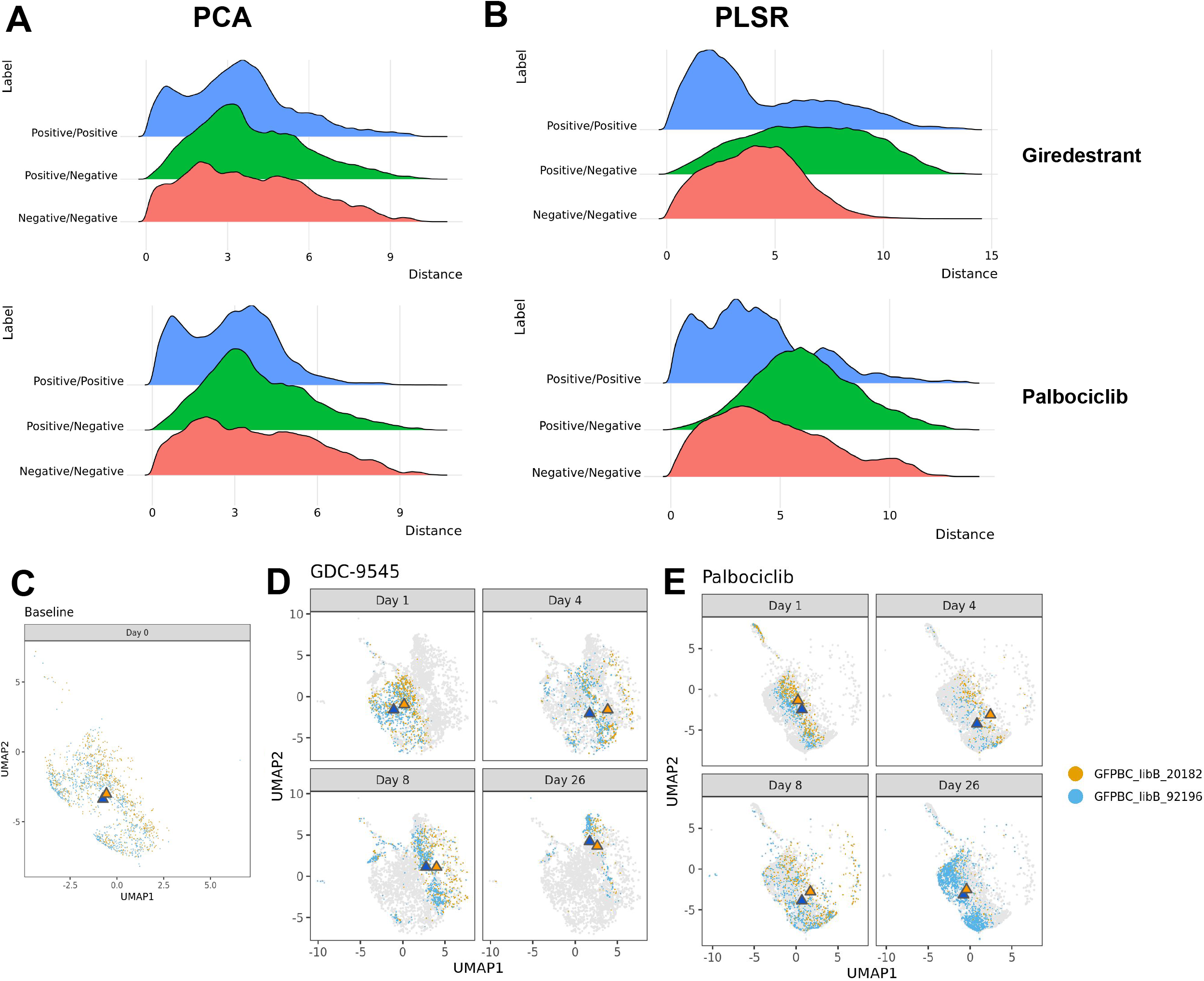
Ridge plots depict the pairwise distance between two PS cells (blue), two NS cells (red), and one PS cell and one NS cell (green) when PCA dimension reduction is used (A). The distance between PS to PS cells and NS to NS cells is reduced, while PS and NS cells are more separated with PLSR (B). The transcriptomes of a NS clone (orange) and PS clone (blue) and their centroids (triangles) are plotted together pre-treatment, showing they are not separated (C). When treated with giredestrant (D) or palbociclib (E) the transcriptomes of these clones change throughout treatment, but the clone centroids remain together and are not separated by treatment.

**Figure S4.**
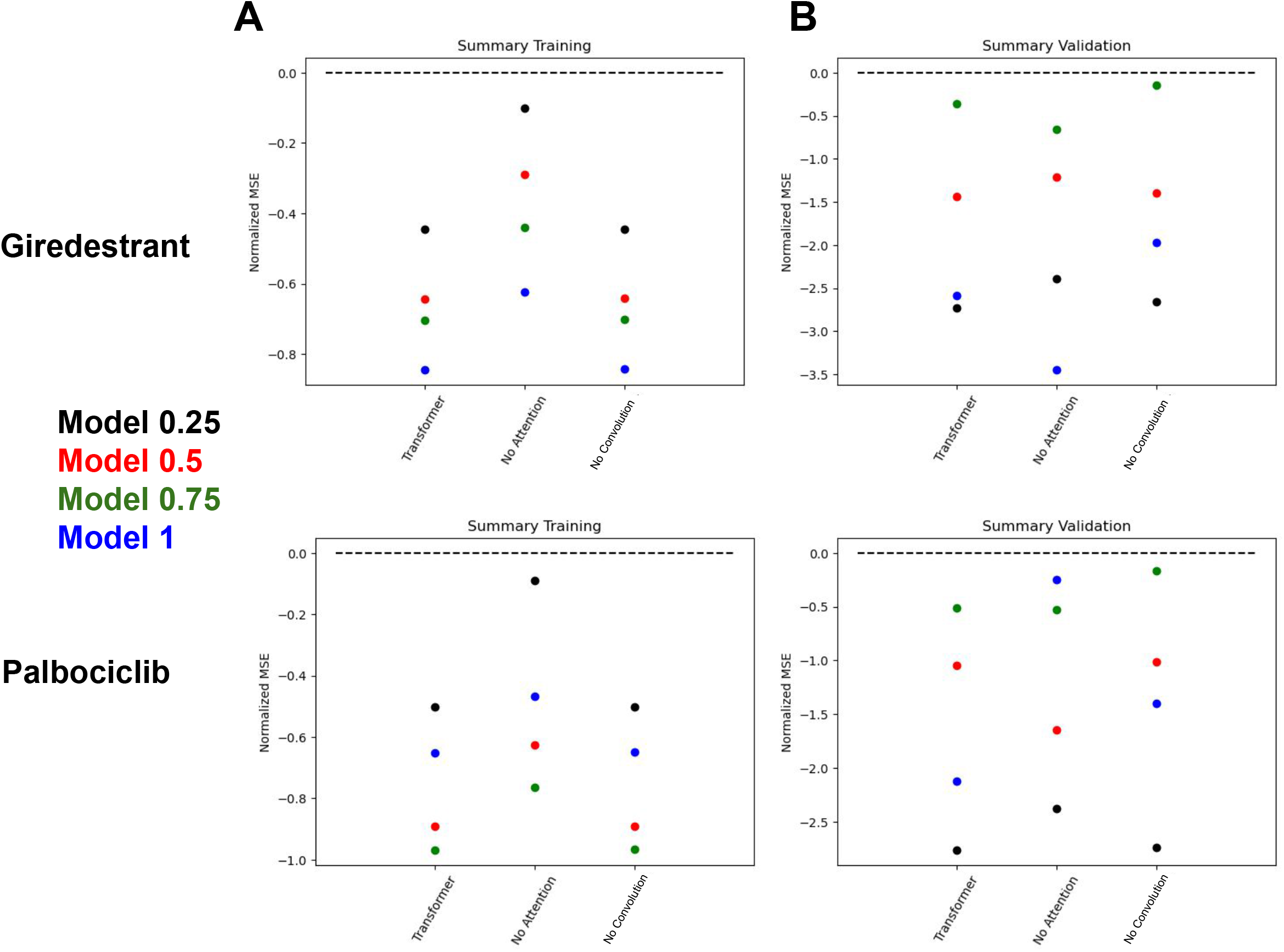
The model error in training is plotted relative to multivariate regression (dashed line) for modeling up till each of the four testing increments at 0.25 (black), 0.5 (red), 0.75 (green), and the full fraction of the clonal data (blue). Error is represented as the mean error for the modeled barcodes for the full TraCSED model (left), TraCSED without the attention layers (middle), and TraCSED without the convolutional layers (right) for both giredestrant and palbociclib (A). The model error is then also plotted in the same way for the validation data (B).

**Figure S5.**
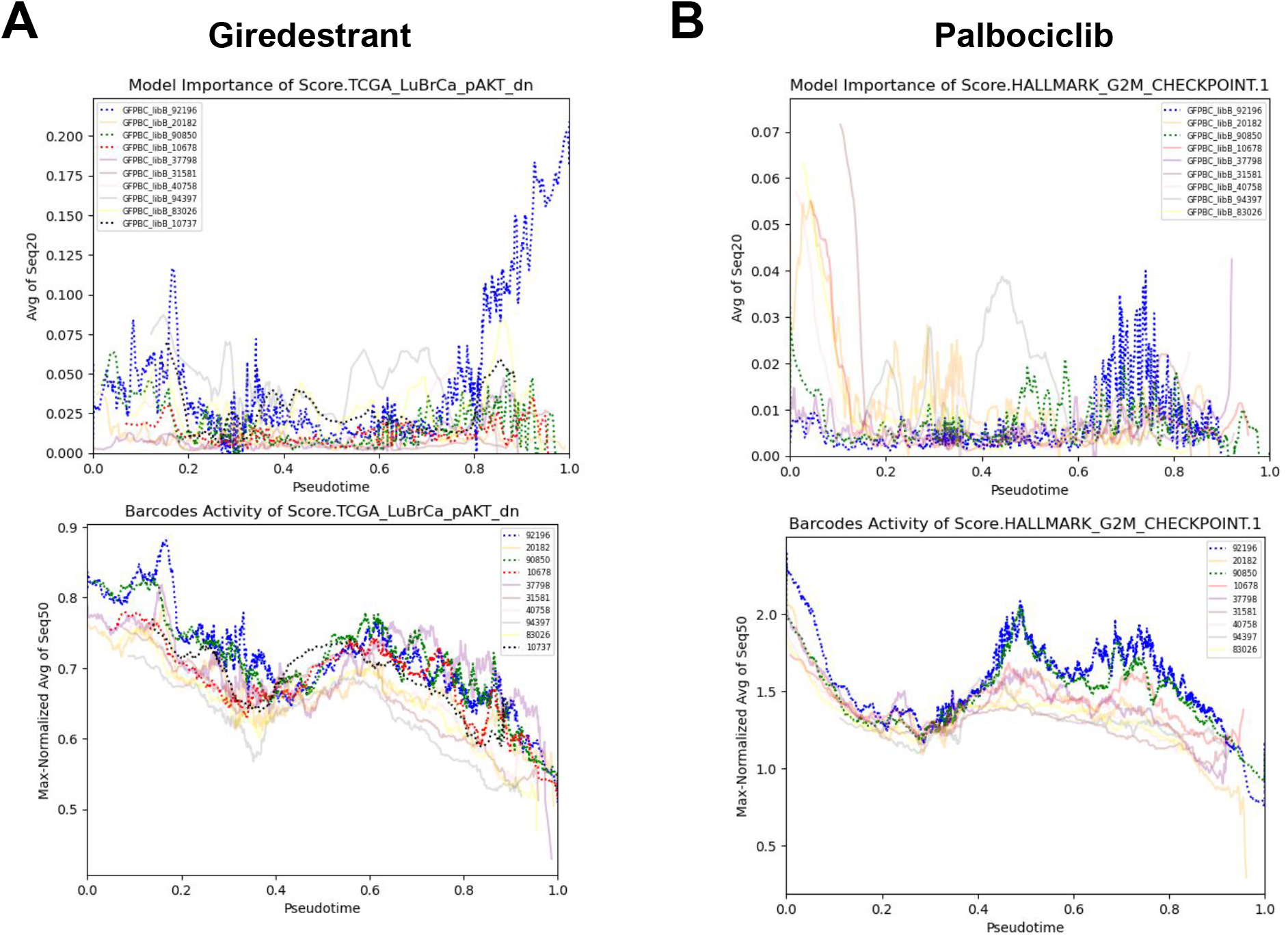
Smoothed model importance (averaging 20 adjacent cells) was plotted for PS clones (bold, dashed lines) and NS clones (faint, solid lines) for the AKT pathway in giredestrant treatment (upper panel). Smoothed pathway activity (averaging 50 adjacent cells) was also plotted (lower panel, A). Similarly, the smoothed model importance (upper panel) and pathway activity (lower panel) were plotted for the G2M checkpoint pathway in palbociclib treatment (B).

**Figure S6.**
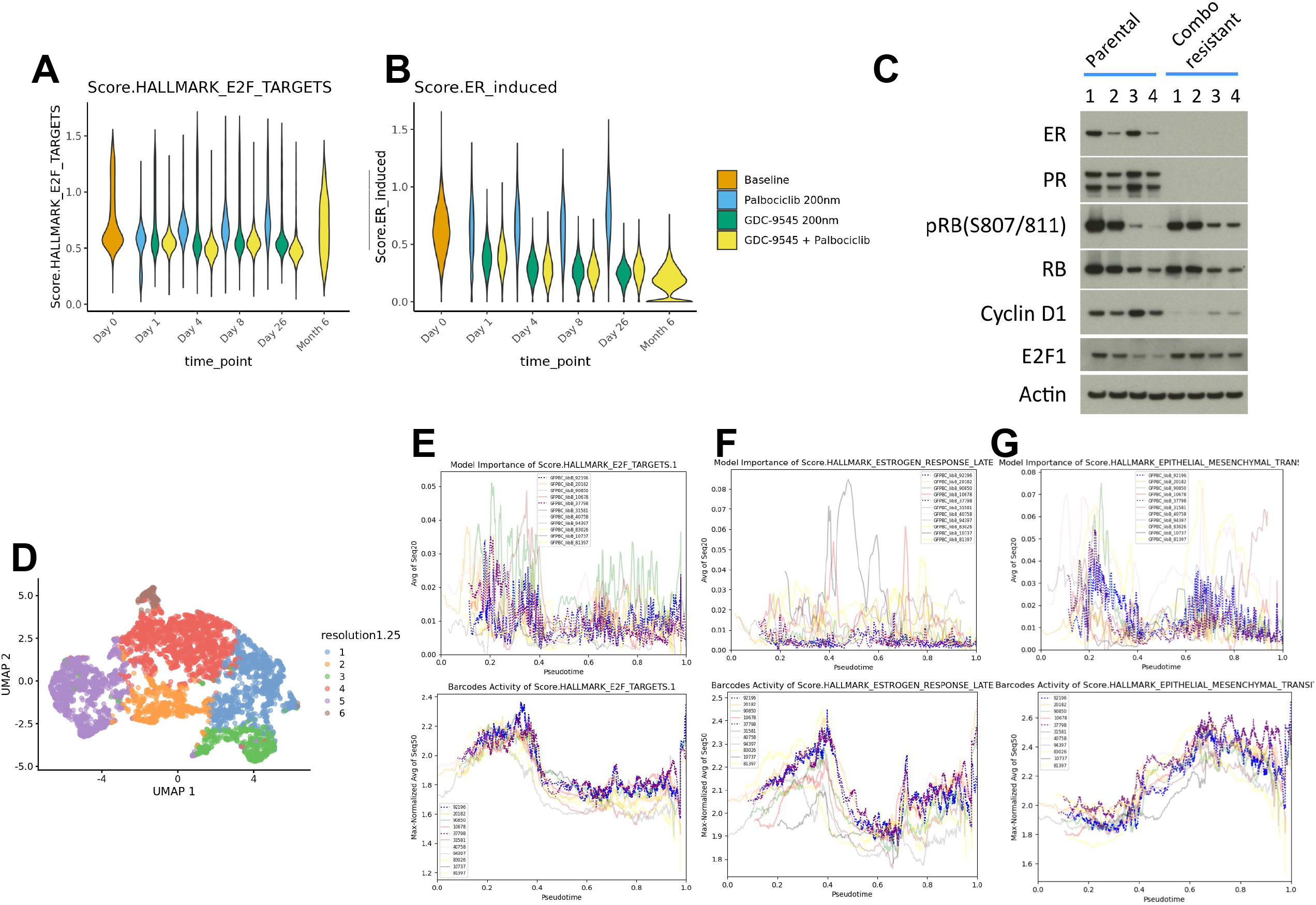
Pathway scores for E2F targets (A) and ER induced activity (B) is plotted across pre-treatment (orange) and for the duration of treatment with palbociclib (blue), giredestrant (green), and the giredestrant-palbociclib combination therapy (yellow). The cluster results from the PLSR approach are plotted for the giredestrant and palbociclib combination treatment (C). Smoothed model importance (averaging 20 adjacent cells) was plotted for PS clones (bold, dashed lines) and NS clones (faint, solid lines)(upper panel). Smoothed pathway activity (averaging 50 adjacent cells) was also plotted (lower panel). This was done for the E2F targets pathway (C), Estrogen Response Late (D), and EMT pathway (E).

**Figure S7.**
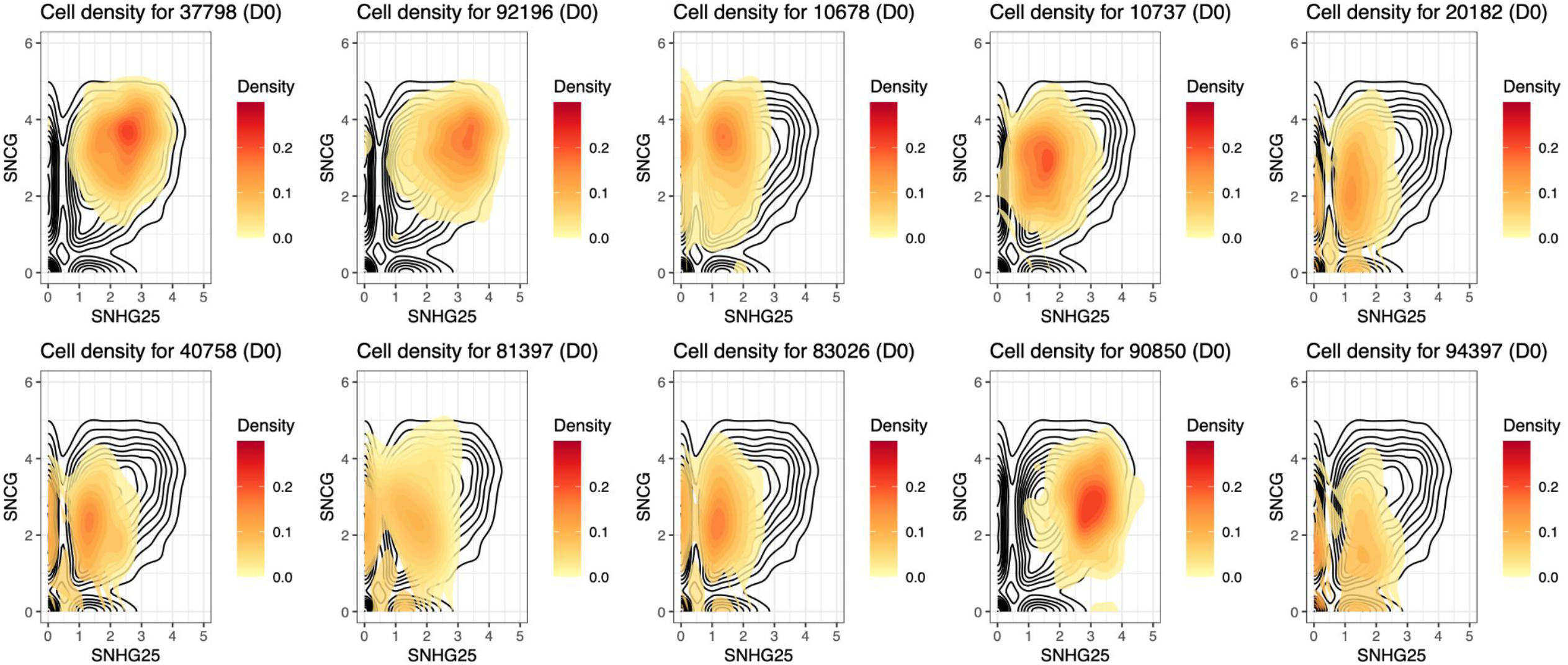
Distribution of cells at baseline based on the expression of SNHG25 and SNCG. Black outline shows all cells; color density shows cells from each clone individually.

